# FcεRI+IgE+ monocytes are linked to atopy and allergic inflammation with distinct phenotypes and enhanced antiviral responses

**DOI:** 10.64898/2026.06.22.733587

**Authors:** Jinyi Wu, Bailey Matthews, Siva Solleti, Regina K. Rowe

**Affiliations:** Department of Pediatrics, University of Rochester Medical Center, Rochester, NY 14642; Biomedical Genetics and Genomics Graduate Program, University of Rochester Medical Center, Rochester, NY 14642

**Author notes:** **Corresponding author:** Regina K. Rowe, MD PhD, University of Rochester Medical Center, 601 Elmwood Ave, Box 690, Rochester, NY 14642.

**Keywords:** IgE, FcεRI, atopy, monocyte, antigen presenting cell, allergic airway disease, asthma, antiviral response

## Abstract

Monocytes are critical regulators of allergic inflammation, whose functions are modified by IgE-driven processes. Monocytes are heterogeneous; comprised of multiple subsets which implies differential functions. In allergic inflammation, this heterogeneity is likely influenced by IgE-mediated effects. We sought to identify phenotypically distinct monocyte subsets related to allergic disease and then further delineate functional differences in cytokine release and antiviral responses. Using high dimensional spectral flow cytometry, we identified monocyte surface phenotypes directly related to surface levels of the high affinity IgE receptor (FcεRI) and surface-bound IgE. FcεRIα+IgE+ monocytes, or FIMs, correlated with allergic disease and the level of atopy (i.e. serum IgE levels) of individual subjects. The FIM population also had differential surface expression of other molecules of monocyte maturation, which closely resembled a type 2 conventional dendritic cell (cDC2) phenotype. Functionally, FIMs had enhanced antiviral responses and IgE-driven IL-10 cytokine release. Finally, we showed that FIMs could be identified at higher levels in lung tissue from individuals with asthma. This study supports that atopic disease drives differential monocyte phenotypes, with the FIM population, specifically, as a more mature cell population closely related to dendritic cells with enhanced antiviral responses. The presence of monocytes in lung tissue during lethal asthma exacerbation further supports a role in regulating tissue inflammatory responses in allergic airway disease.

## Introduction

Monocytes are innate cells that comprise 10-20% of the circulating mononuclear leukocytes in the peripheral blood, and are critical players in the innate immune responses. During inflammation, monocytes migrate from blood into tissues, regulating inflammatory responses through cytokine and chemokine release and phagocytosis. Their actions then “bridge” innate and adaptive immune responses through antigen presentation and T cell stimulation. Monocytes, however, are heterogeneous suggesting specific functional subsets, which are likely influenced in disease states. While most studies identify monocyte subsets based on CD14 and CD16 expression (e.g. classical, intermediate, and non-classical), recent studies using single cell transcriptomic approaches have identified phenotypes and functions beyond these two canonical markers ^1^.

IgE-driven allergic inflammation is known to impact monocyte functions. Activation of the high affinity IgE receptor, FcɛRI, via IgE-crosslinking, has significant effects on cellular maturation, cytokine and chemokine release, antiviral responses, and T cell differentiation ^2–4^. Serum IgE levels correlate with expression of FcɛRI on monocytes, which also correlates with the severity of allergic disease ^5,6,7.^ It has previously been shown that a population of CD14+ monocytes positive for FcɛRI also co-express CD2, with this population acquiring dendritic cell (DC)-like activities ^7^. Similarly, recent single cell sequencing analyses support that multiple monocyte populations exist which are transcriptionally distinct^1^ and can be identified in allergic airways^8^. While these studies suggest differential monocyte functions in the setting of allergic disease, how IgE levels directly regulate monocyte development, phenotypic diversity, and function is not currently well understood.

Monocytes are recruited to the airways in allergic asthma^8^, including virus-induced exacerbations ^9,10^. Monocytes in the lungs also correlate with respiratory virus infection severity ^11^. In allergic asthma, airway proinflammatory and antiviral cytokine, and T cell responses have been shown to correlate with monocyte and macrophage gene and pathway signatures ^8,12^. Changes to monocyte phenotypes in the peripheral blood by systemic allergic inflammation may dictate how these cells function upon tissue migration. Thus, understanding the mechanisms by which IgE-driven allergic disease modifies monocyte phenotypes and functions – both in the blood and tissue - is crucial to understanding the role of monocytes in allergic disease. This study aimed understand how atopy impacts monocyte phenotypic diversity and functions. We show that IgE-mediated allergic disease is related to a population of atopic monocytes which express high levels of FcεRIα and have high surface-bound IgE (FcεRIα+IgE+ monocytes or FIMs). FIMs also co-express other cell surface molecules that resemble type 2 conventional DCs (cDC2) and have enhanced cytokine and antiviral maturation responses. FIMs were also identified in lung tissue of lethal asthma exacerbations. These data support that atopic disease promotes the development of monocyte subsets with differential functions which may have roles in regulating tissue inflammation.

## Results

### FcεRIa+IgE+ monocytes (FIMs) are related to allergic disease and serum IgE levels

While monocytes from individuals with atopy have higher surface expression of the high affinity IgE receptor (FcεRI) ^6,13^, it was not known how this related to surface bound IgE and expression of other (non-atopic) surface molecules. To test this, we recruited a cohort of adults and children with and without allergic disease, and a range of serum IgE levels (Table 1). Since allergic history was by self-report, we confirmed that allergic subjects had significantly higher levels of serum IgE (Fig. 1A). This allowed us to compare measurements by either allergic history or serum IgE levels (high >100IU/mL vs. low <100IU/ml ^14^). We performed flow cytometry on PBMCs using a high parameter spectral flow cytometry panel, with 20 surface antigens comprised of cell type specific markers, markers of atopy, and molecules associated with monocyte functions (*e.g.* antigen presentation, T cell stimulation, cellular maturation) (Supp. Table 1). Neither allergic disease or the level of atopy (*i.e.* serum IgE levels) were related to proportions of the major monocyte subsets (CD14+ or CD14+CD16+) (Supp. Fig. 1, Supp. Fig. 2). We then evaluated the CD14+ monocyte population based on surface levels of FcεRIα and bound IgE. While the majority of CD14+ monocytes expressed FcεRIα, there was a notable proportion with high levels of surface-bound IgE (FcεRIα+IgE+) (Fig. 1B). These FcεRIα+IgE+ monocytes, or FIMs, were higher in allergic subjects, (p=0.095) (Fig. 1C). Inspection of individual subject data identified 2 outliers in the non-allergic group with a high % of FIMs who also had high serum IgE (126 and 318 IU/mL) levels, indicating that serum IgE may correlate with the FIM+ population. When comparing based on IgE levels, subjects with high serum IgE had a significantly higher proportion of FIMs, regardless of allergic history (Fig. 1C). We wanted to determine whether this was specific to the high affinity IgE receptor, FcεRI, or if all IgE receptors were increased, as monocytes also express the low affinity IgE receptor, CD23. However, there was no relationship between either serum IgE levels or allergic history and CD23hi monocytes (Supp Fig. 2). This supports specificity of serum IgE in promoting development of the FcεRIα+IgE+ monocyte (FIM) population and not simply increased expression of all IgE receptors.

**Figure 1:**
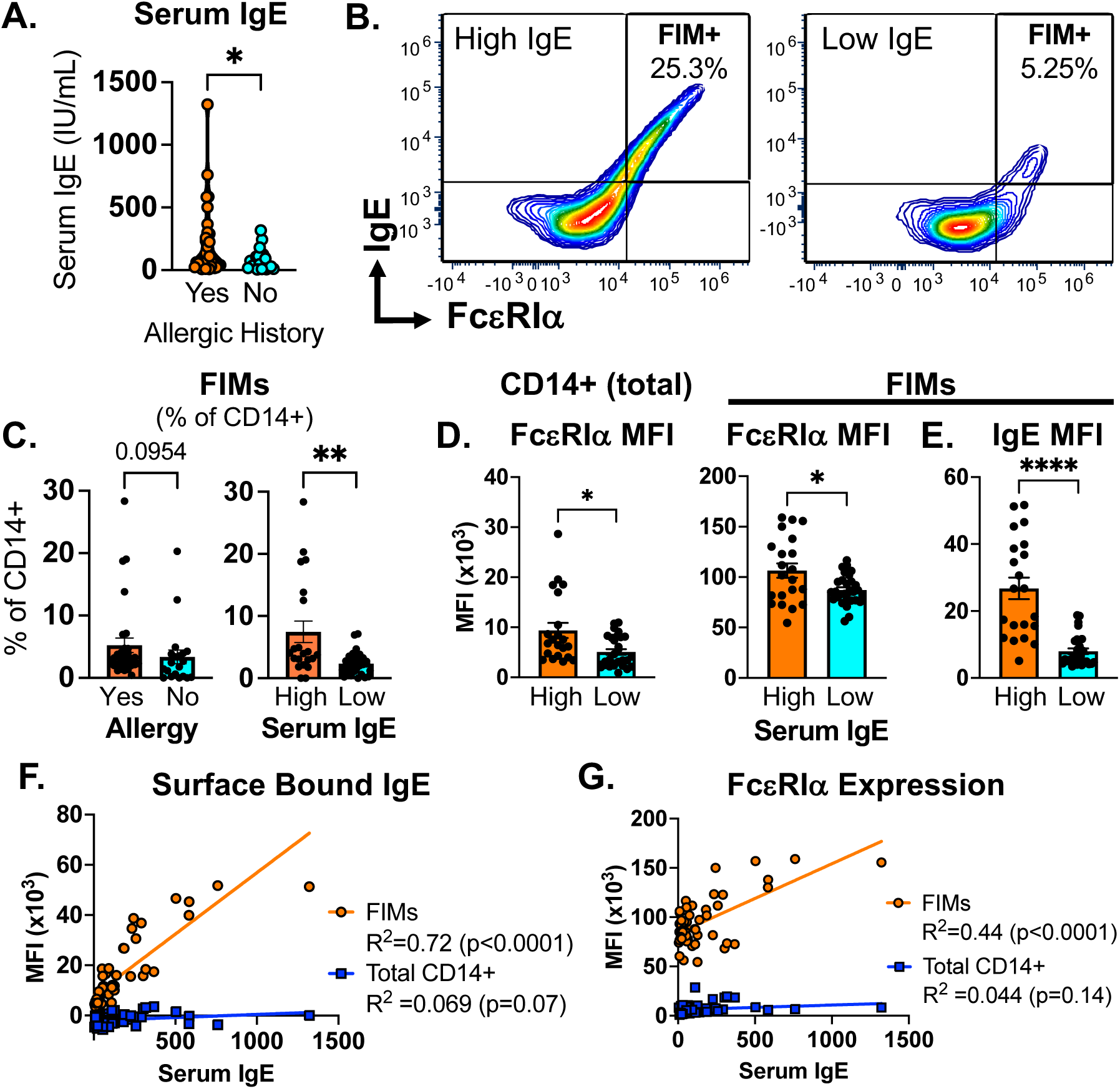
IgE-mediated allergic disease is related to FcεRIα+IgE+ (FIM) subset. PBMCs from allergic and non-allergic subjects were analyzed by spectral flow cytometry to identify monocyte and dendritic cell subsets. (A) Serum IgE levels between subjects with (yes) and without (no) allergic history. (B) Flow cytometry of the FIM+ population in subjects with either high or low serum IgE levels. (C) Allergic and high IgE subjects have greater % of the FIM+ population. (D) FcεRIα surface expression on total CD14+ and FIM+ populations, and (E) surface-bound IgE levels were compared for the FIM+ population between high and low serum IgE subjects. (F-G) Linear regression analaysis of (F) surface IgE and (G) FcεRIα levels with serum IgE levels for both CD14+ and FIM+ monocytes.

**Table 1:** Subject Demographics.

| Table 1: Subject Demographics |  |  |  |  |  |  |
| --- | --- | --- | --- | --- | --- | --- |
| Number (n) | Total (n=50) |  | Child (n=11) |  | Adult (n=39) |  |
| Age (y; mean, range) | 25 (5-57) |  | 8 (5-11) |  | 30 (19-57) |  |
| Sex (Female/Male) | F (30); M (20) |  | F (6); M (5) |  | F (24); M (15) |  |
| Serum IgE IU/mL mean (range; SD) | 162<br>(5-1322; +/-239) |  | 235<br>(8-1322; +/-384) |  | 141<br>(5-760; +/-181) |  |
| High (>100IU/mL)<br>Low (<100IU/mL) | High (21); Low (29) |  | High (6); Low (5) |  | High (15); Low (24) |  |
| Allergic History (n) | No (20) | Yes (30) | No (6) | Yes (5) | No (14) | Yes (25) |
| Serum IgE mean (range SD) | 82<br>(5-318; +/-85) | 215<br>(9-1322; +/-291) | 85<br>(8-126; +/-48) | 415<br>(22-1322 +/-540) | 80.6<br>(5-318; +/-99) | 175<br>(9-760; +/-209) |
| Diagnosis (n) | Allergic Asthma (11), Allergic Rhinitis (21), Atopic Dermatitis (7), Food Allergy (11), Medication Allergy (6), Chronic Sinusitis (1) |  |  |  |  |  |

We then evaluated the relationship between serum IgE and surface levels of FcεRIα and IgE, comparing the total CD14+, CD23hi, and FIM populations. Subjects with high serum IgE had uniformly higher FcεRIα surface levels in all three monocyte populations (Fig. 1D, Supp.Fig. 2). However, surface-bound IgE showed a divergent relationship. Only FIMs had increased surface-bound IgE when comparing subjects with high versus low serum IgE (Fig. 1E). Across all subjects, serum IgE positively correlated with surface-bound IgE levels in the FIM population, however no correlation found in the total CD14+ population (Fig. 1F). Similarly, FcεRIα expression only correlated with serum IgE in the FIM population (Fig. 1G). In contrast, CD23 surface expression was not increased in high IgE subjects (Supp. Fig. 2). Instead, subjects with low serum IgE had a small population of monocytes (mean 1% of CD14+), which had higher CD23 levels with detectable surface-bound IgE but low FcεRIα expression (FcεRIα-IgE+ Supp. Fig. 2). This supports differential IgE binding across monocyte subsets in relationship to the IgE receptors, and that serum IgE levels do not equally regulate IgE binding and IgE receptor expression across the monocyte population uniformly.

### FIMs monocytes have a distinct cell surface phenotype similar to cDC2s

Additional molecules involved in monocyte maturation and T cell regulation (Supp. Table 1) but not associated with atopy were also measured from the flow cytometry analysis. From this data, we identified other surface phenotypes associated with the FIM population. When compared to total CD14+ monocytes, many molecules, in addition to IgE and FcεRIα, were increased on the surface of FIMs including MHC II (HLA-DR), CD123 (IL3Ra), CD1c (BDCA-1), and ILT-3 (Fig. 2A-H). However, multiple proteins were lower, such as CD23, CD14, CD86, CD40, ICAM-1, ILT-4, and Siglec-6 (Fig. 2I-P). Several proteins were also related to serum IgE levels. For example, subjects with high IgE had increased CD14 and ILT-4 on FIMs (Fig. 3A-B), despite lower overall surface expression on this population compared to total monocytes. CD1c, while higher in FIMs overall, was increased in low serum IgE subjects (Fig. 3C). Siglec-6, similarly showed higher levels in low IgE subjects but in both the CD14+ and FIM populations (Fig. 3D). This data suggests that IgE is related to differential expression of molecules not typically associated with atopy, and a complexity in how monocyte phenotypes differ across individual subjects and levels of atopy.

**Figure 2:**
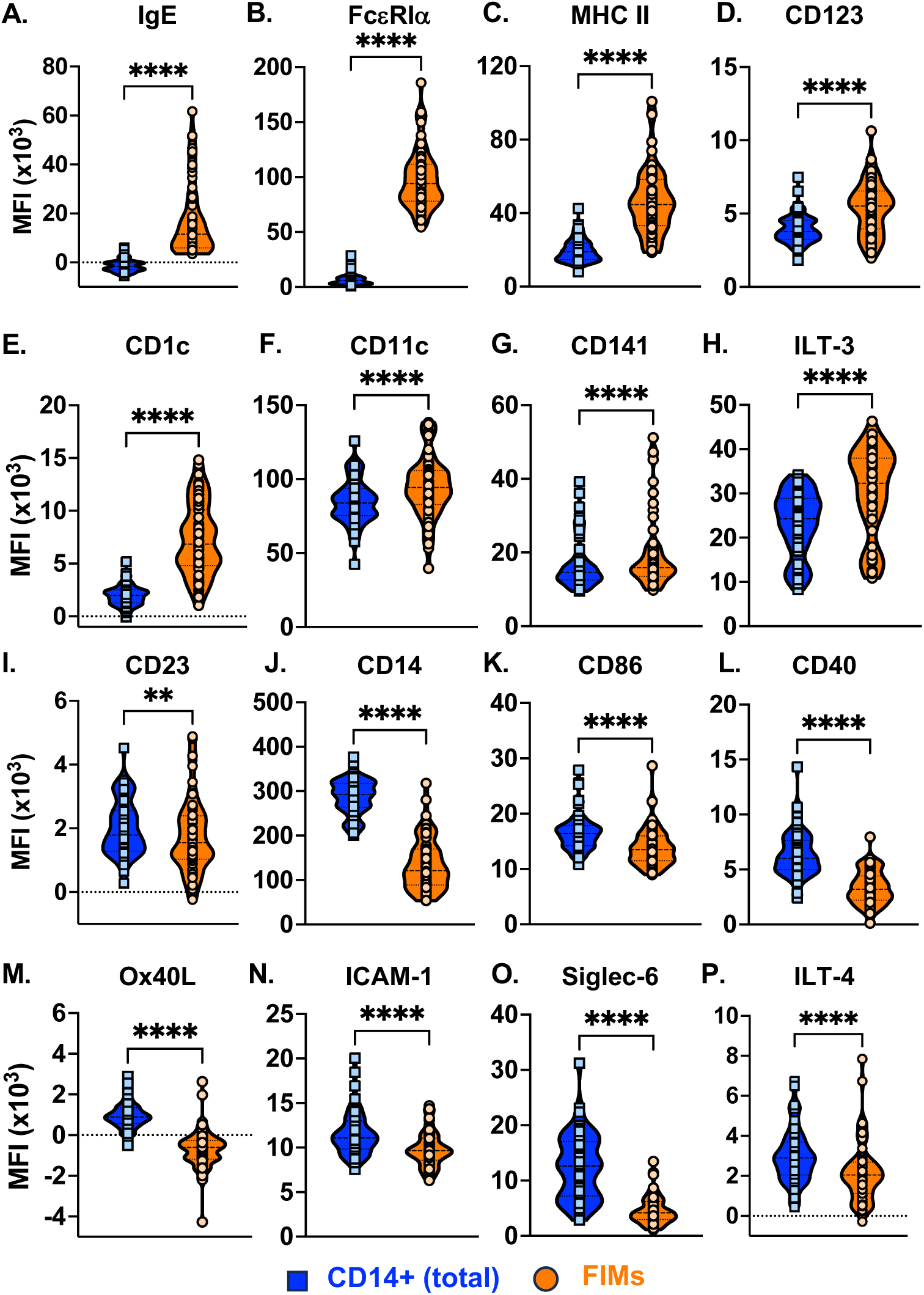
FIMs have a distinct cell surface phenotype. Spectral flow cytometry panel that included 19 cell surface markers to phenotype the total CD14+ population (blue) and the FIM (orange) subset. Shown are 16 proteins (A-P) which demonstrated significant differences between the total CD14+ monocyte population and FIM population where protein was either increased (A-H) or decreased (I-P) on FIMs compared to the total CD14+ population. Paired two-tailed t-test. *p<0.05, **p<0.01, ***p<0.001, ****p<0.0001.

**Figure 3:**
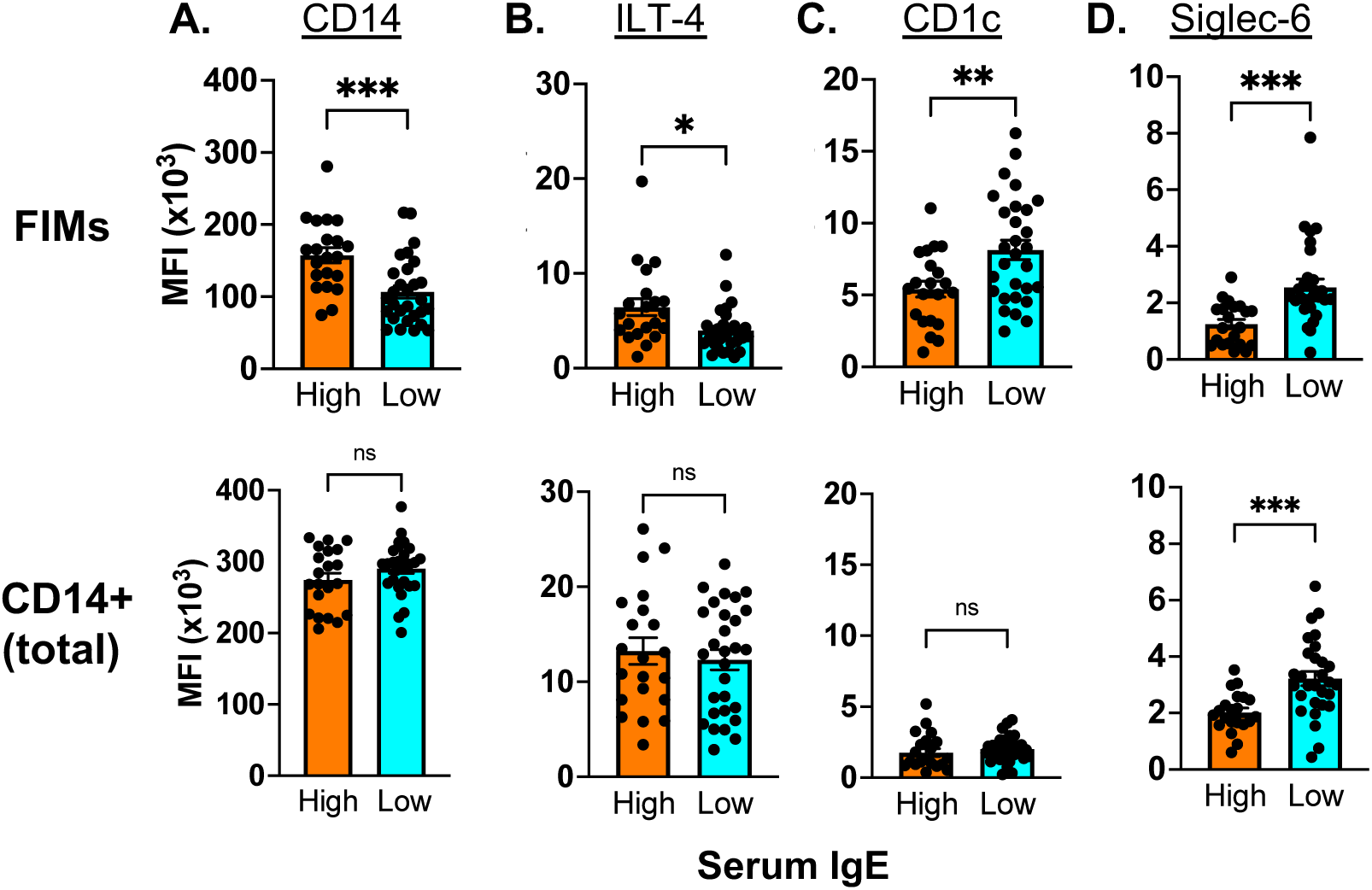
Serum IgE is related to surface phenotypes in FIMs and CD14+ monocytes. Surface protein levels for (A) CD14, (B) CD1c, (C) ILT-4, and (D) Siglec-6 were compared between subjects with high versus low serum IgE levels on FIMs (top row) and the total CD14+ monocyte (bottom row) populations. Unpaired t-test with Welch’s correction; *p<0.05, **p<0.01, ***p<0.001, and not significant (ns).

Using dimensionality reduction methods UMAP and PHATE, we evaluated how FIMs were related to other monocyte and dendritic cell populations. Combining both canonical gating strategies (Fig. 1 and Supp. Fig. 1) and UMAP dimensional reduction, we identified clusters representing all major monocyte and DC populations, including classical (CD14+CD16-), intermediate (CD14+CD16+), and non-classical (CD14^low^CD16+) monocytes, plasmacytoid DCs (pDCs), conventional (myeloid) dendritic cells (cDC1 and cDC2), and basophils (Fig. 4A). FIMs were identifies as a population off the classical monocyte cluster (Fig. 4A; Population 1 vs. 4), however were distinct from the main populations of CD14+ or CD16+ monocytes (Fig. 4B). UMAP combined with individual surface molecule expression patterns (*e.g.* increased MHC II and CD1c; Fig. 2), suggested that the FIMs were closely related to CD1c+ DCs or cDC2s (Fig. 4A, Populations 4 vs. 7). Cellular trajectories were then analyzed across monocyte and DC populations using PHATE, which preserves both local and global structure of the data ^15^. This suggested a continuum across FcεRIα and IgE levels in monocytes with FcεRIα-/low monocytes closely related to CD16+ monocytes and cells moving along a trajectory (Fig. 4C) with decreasing in CD14 expression, while increasing in FcεRIα, MHC II, and bound IgE (Fig. 4D). FIMs were adjacent to the cDC2 population, however weren’t simply CD14+ DCs as expression of common DC markers were distinct between FIMs and CD1c+ DCs (Fig. 4E).

**Figure 4:**
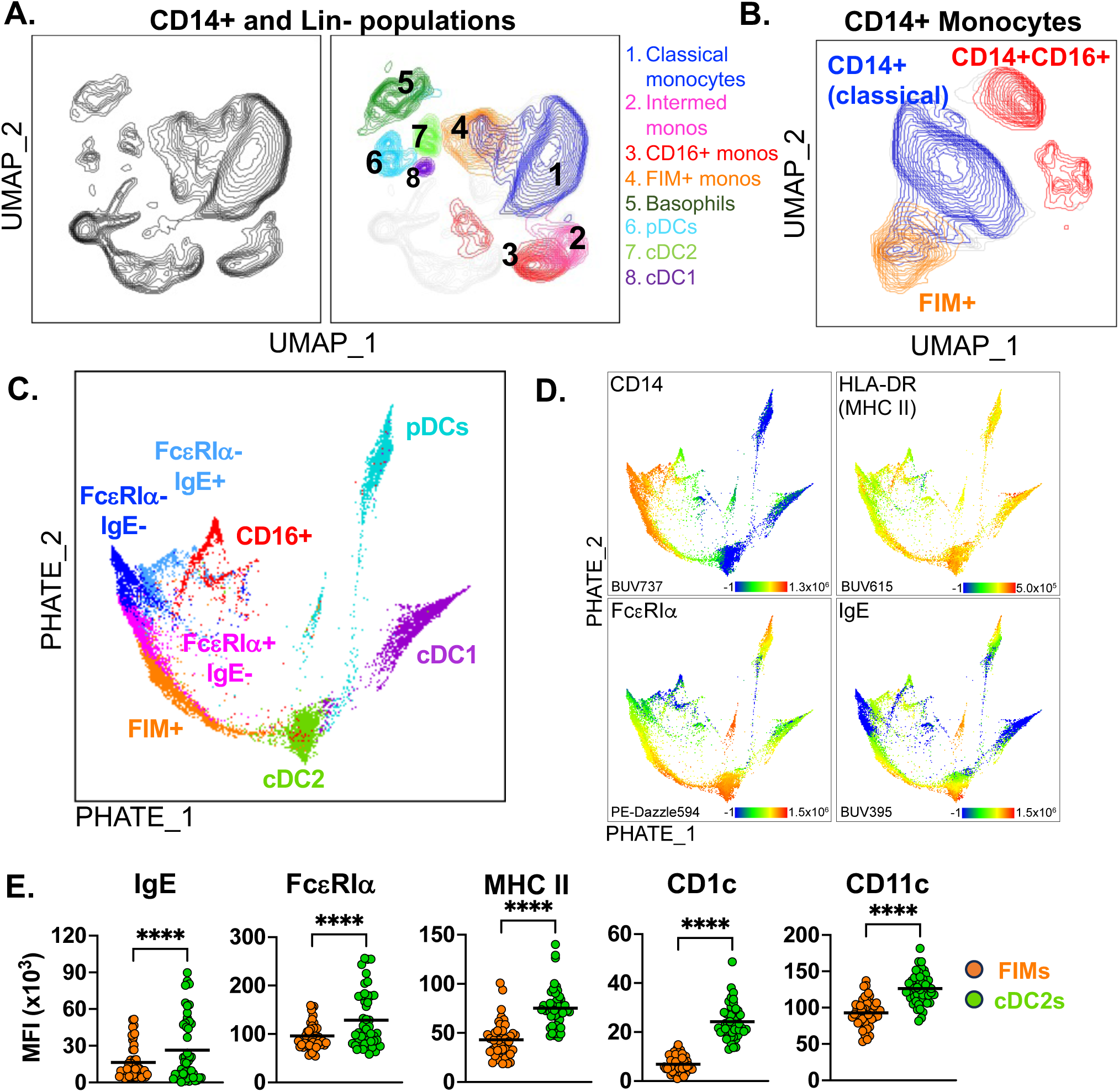
FIMs are phenotypically similar to cDC2s. Dimensionality reduction by UMAP with overlay of cell populations for (A) monocyte, DC, and basophils and (B) CD14+ populations. (C) PHATE trajectory analysis of monocyte and DCs with (D) heatmap of selected surface markers. (E) Comparison of FIM and cDC2 surface levels of select surface markers; paired t-test, ****p<0.0001.

### FIMs have differential antiviral and IgE-driven responses

We previously showed that IgE-mediated stimulation inhibits monocyte antiviral responses, including virus-induced maturation and antiviral CXCL10 (IP-10) release, while resulting in IL-10 production ^2–4^. We first wanted to determine if allergic disease impacts these IgE-specific antiviral phenotypes. Human monocytes were isolated from a subset of allergic (n=12) and non-allergic (n=9) groups. Total monocytes were stimulated with liposomal-poly(I:C) (lipo-pIC) with and without IgE-crosslinking (αIgE), and IP-10 and IL-10 release were evaluated (Fig. 5A-B). The allergic group had decreased lipo-pIC induced IP-10 release (Fig. 5A), and increased IL-10 release in the setting of IgE-crosslinking and lipo-pIC stimulation (Fig. 5B).

**Figure 5:**
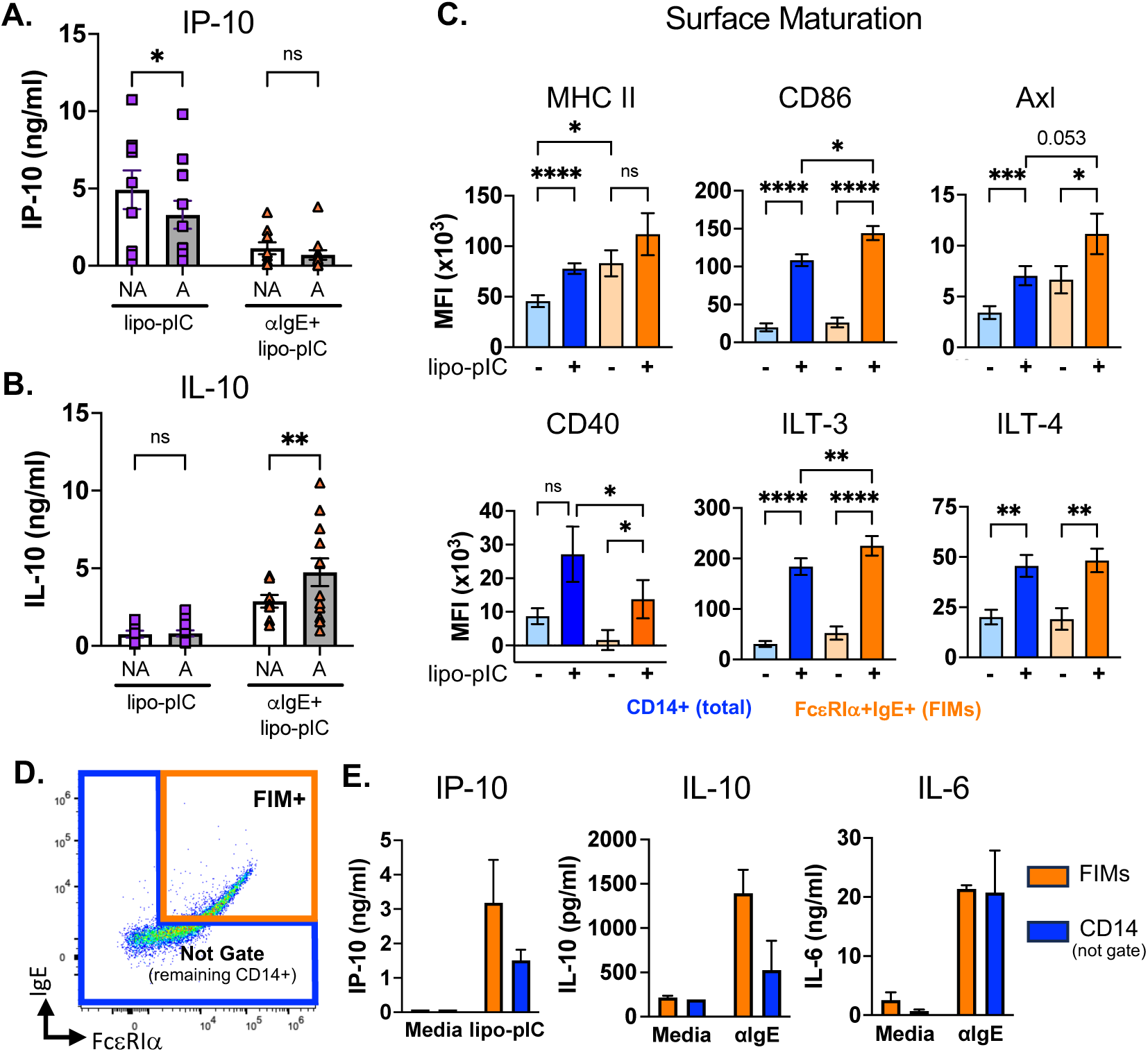
Monocyte maturation and cytokine responses are affected by IgE-driven allergic disease. (A-B) Monocytes were isolated from allergic (A; n=12) and non-allergic (NA; n=9) subjects and stimulated with liposomal-poly(IC) (lipo-pIC) with or without IgE crosslinking (αIgE). Supernatants were analyzed at 24 hours by ELISA for (A) IP-10 and (B) IL-10 levels. Two-way ANOVA with multiple comparisons; ns = not significant; *p<0.05, **p<0.01. (C) CD14+ monocytes were isolated, stimulated with lipo-pIC for 24 hours, and evaluated by flow cytometry for surface expression of MHC II, CD86, Axl, CD40, ILT-3, and ILT-4. Expression was compared between the CD14+ total monocyte population and FIMs. n=8 subjects analyzed by repeated measures ANOVA; ns= not significant; *p<0.05, **p<0.01, ***p<0.001, ****p<0.0001. (D) Monocytes from 3 subjects with high % of FIMs were sorted for FIM+ or remaining (Not Gate) CD14+ populations. (E) Sorted populations were stimulated with either media alone, lipo-pIC, or IgE crosslinking (αIgE). Supernatants were analyzed at 24 hours for IP-10, IL-10, and IL-6 by ELISA.

We then compared responses between the FIM+ and CD14+ populations. Many of the proteins differentially expressed in the FIM population are associated with antiviral responses and cellular maturation. Isolated CD14+ monocytes (n=8 subjects) were stimulated with lipo-pIC for 24 hours and evaluated by flow cytometry (Fig. 5C). Cell surface protein expression was compared between the total CD14+ and FIM+ populations. Lipo-pIC stimulation increased surface expression for MHC II, CD86, Axl, CD40, ILT-3, and ILT-4 (Fig. 5C), with differences noted between the total CD14+ population and FIMs. MHC II upregulation did not reach significance in the FIM population as compared to CD14+ monocytes, likely due to higher baseline expression. However, other markers including CD86, Axl, and ILT-3, were upregulated to a greater extent on FIMs compared to total CD14 monocytes. CD40, conversely was upregulated less in FIMs, while the ILT-4 showed equal upregulation in both populations (Fig. 5C). To compare cytokine and chemokine release, we sorted FIMs from the remaining CD14+ population (Fig. 5D). Equal cell concentrations (cells/mL) were stimulated with either lipo-pIC or αIgE, and IP-10, IL-10, and IL-6 were measured by ELISA (Fig. 5E). The FIM population trended towards higher levels of IP-10 and IL-10 production, while IL-6 was similar for both populations (Fig. 5E). This suggested enhanced antiviral and IgE-driven responses by FIMs consistent with a more mature functional phenotype, in line with the surface phenotyping analysis.

### Monocyte phenotypes differ in lung tissue of subjects with asthma

Since monocytes are recruited from the peripheral blood into tissue, we determined whether FIMs were also present in lung tissue. Flow cytometry was performed on dissociated lung tissue from post-mortem samples from subjects with (n=7) and without (n=7) history of asthma (Supp. Table 2), with acute asthma exacerbation as the cause of death in six subjects. Anti-CD45 was added to the flow cytometry panel (Supp. Table 1) to identify hematopoietic cells (Fig. 7A) from the heterogenous mixture. Lung tissue from subjects with asthma did have more CD45+ cells present (data not shown), consistent with inflammation. Although there was no difference in the proportion of total CD14+ monocytes, asthma lung tissue did have a higher % of FIMs (Fig. 6A-B). UMAP analysis of CD14+ monocytes showed differences between asthma and non-asthma lung tissue (Fig. 6C, Supp. Fig. 3). Evaluation of specific surface proteins between CD14+ and FIM+ populations, showed many similarities with blood monocytes including lower CD14 and higher MHC II, ILT-3, CD123 (Fig. 6D) on FIMs. Other proteins, however, were opposite to observations in blood monocytes showing higher expression of CD23, CD86, CD80, ICAM-1, Axl, ILT-4, Siglec-6, and CD16 on FIMs compared to the total monocyte population (Fig. 6E).

**Figure 6:**
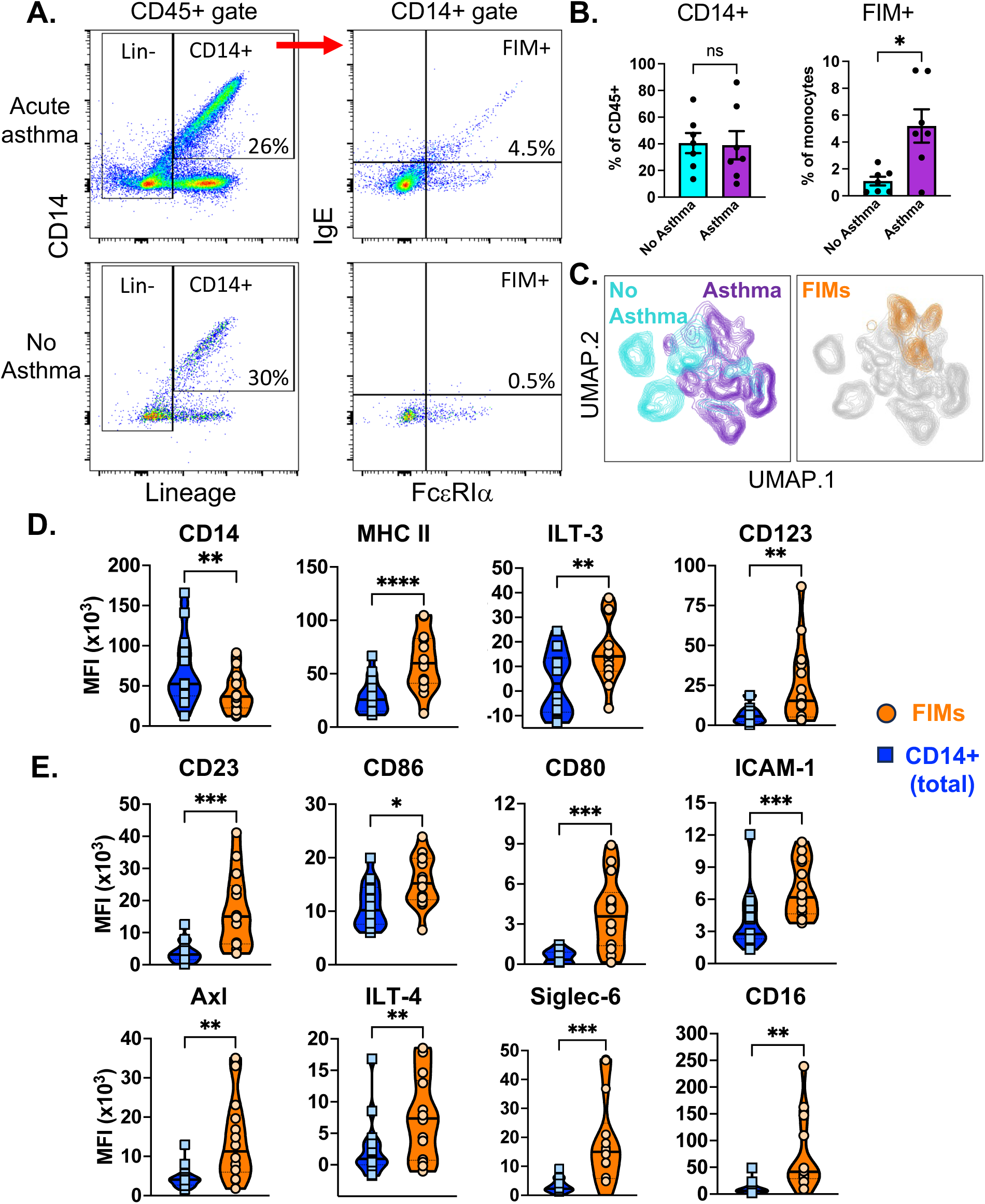
FIMs are present in lung tissue of subjects with asthma. Dissociated lung tissue from post-mortem donors (n=14) were analyzed by flow cytometry for CD14+ and FIM populations. (A) Flow cytometry gating for lungs from a representative sample with acute asthma (top) and no asthma (bottom) history. (B) Comparison of total CD14+ and FIM+ populations in no asthma (n=7) vs. asthma (n=7) lung tissue. Mann-Whitney test; ns=not significant; *p<0.05. (C) Dimensionality reduction by UMAP with overlay of no asthma (cyan) versus asthma (purple) subjects and the FIM population (orange). (D) Comparison of select markers which showed significant differences between the CD14+ monocyte population and the FcεRIα+IgE+ monocyte population in lung tissue. Paired two-tailed t-test; *p<0.05, **p<0.01, ***p<0.001, ****p<0.0001.

Four of these markers, CD23, CD123, Siglec-6, and ILT-3, were also significantly increased on the total CD14+ monocyte population when comparing asthma to non-asthma tissue (Supp. Fig. 4), consistent with an overall upregulation in allergic inflammation.

## Discussion

This study identified a strong relationship between IgE-driven allergic disease and monocyte surface marker and functional phenotypes. Prior to this study, less was known about how atopic disease modifies different monocyte populations. Atopic disease severity is known to correlate with serum IgE levels ^16–19^ and increased FcεRI expression on mast cells, basophils, and dendritic cells ^13,20–23^. However, in monocytes, serum IgE and FcεRI receptor expression have shown less of a relationship ^24–26^, suggesting different regulatory mechanisms. Similar to previous studies ^7,24,25^, we identified varying levels of the high affinity IgE receptor, FcεRIα. However, different from other studies, we evaluated the relationship with surface-bound IgE and other proteins not typically associated with atopy. Our data supports that IgE levels and atopy differentially influence monocyte populations beyond just regulation of FcεRI expression and likely influences the overall phenotype – and potentially differentiation – of monocyte subsets.

While a proportion of monocytes in both allergic and non-allergic subjects highly expressed FcεRI, allergic subjects had a subset with both high receptor and surface-bound IgE or FIMs (FcεRIα+IgE+ monocytes). This population was also readily identified and significantly increased in post-morten lung tissue from individuals with asthma. FIMs appear to be an atopy-driven subset directly related to levels serum IgE and allergic disease. While we didn’t formally evaluate IgE-receptor binding, FIMs likely represent a population with high levels of occupied FcεRI receptor which could be readily activated by antigen/allergen binding. Interestingly, in contrast to other reports ^6^, the positive relationship between allergic disease and high serum IgE levels did not exist for the low affinity IgE receptor, CD23. This indicates that IgE does not regulate surface expression of the low affinity and high affinity IgE receptors equally in monocytes. This observed differential IgE-bound phenotype supports that *in vivo*, IgE also does not readily bind to all monocytes expressing the high affinity IgE receptor. Instead, FIMs represent a subset of monocytes influenced by serum IgE, with a greater capacity to upregulate the high-affinity IgE receptor and subsequently bind monomeric IgE.

Previous work by us and others, has shown that IgE-mediated stimulation alters monocyte surface phenotypes and functions, including cellular maturation and cytokine and chemokine secretion. We hypothesized that within the CD14+ monocyte population, the FIM population represents a subset with a different surface phenotype, and differential antiviral and IgE-driven responses. Previous studies have primarily focused on a limited number of surface molecules, such as CD2 and FcεRIα co-expression ^7,24^. High dimensional flow cytometry methods allowed us to perform detailed phenotyping to better define surface phenotypes and incorporate markers not typically associated with allergic disease. We found that FIMs acquire a distinct cell surface phenotype from either the classical CD14+ or non-classical CD16+ monocyte populations. CD14 for example has been shown to be upregulated by IgE-crosslinking ^27^. However, while this molecule may be regulated by IgE-driven processes, surface CD14 expression was actually lower on the FIM population in both blood and lung tissue. We interpret this lower CD14 and higher MHC II expression consistent with cellular maturation, moving towards a monocyte-derived DC phenotype. The UMAP and PHATE dimensionality reduction analyses comparing monocyte and DC populations supports this concept. High expression of MHC II, CD123, and CD1c, are often used as markers of dendritic cells and were all increased on FIMs. PHATE analysis further confirmed that FIMS are most closely related to cDC2s (CD1c+ DCs). FIMs also had increased antiviral-induced surface maturation compared to the remaining CD14+ monocyte population further supporting that this population may possess monocyte-derived DC features. We hypothesize that this population may have increased antigen-presenting functions, and future studies will be needed to further evaluate the antigen presenting and T cell activating capacity of these cells. While it was previously known that monocytes express a range of FcεRI on their surface, how this molecule is related to other cell surface phenotypes was not previously understood. Our data support that IgE-driven allergic disease drives development and differentiation of a specific subset of monocytes, which are present in lung tissue during fatal asthma exacerbations.

Allergic disease has been shown to alter multiple monocyte functions. We showed that monocytes from subjects with allergic disease have decreased antiviral IP-10 (CXCL10) release. IgE-driven activation has been shown to modify multiple functional outcomes of human monocytes. IgE-mediated stimulation triggers release of multiple cytokines, including IL-10, IL-6, IL-1β, TNFα, and IL-23 ^2,3^. IgE-crosslinking also inhibits virus-induced maturation, which alters monocyte-driven CD4 T cell priming and differentiation ^3,4^. We recently showed that stimulation of monocytes with the viral agonist liposomal-pIC combined with IgE crosslinking enhanced IgE-induced IL-10 production ^2^. Our current study demonstrated that this lipo-pIC-driven enhancement of IL-10 is greater in individuals with allergic disease. This supports that IgE-mediated allergic disease promotes a synergistic relationship between antiviral pathway activation and IgE-driven pathways. Most studies have been performed on bulk CD14+ monocytes, thus limiting assessment of functional differences across monocyte subsets. We hypothesized that functional differences were related to surface-bound IgE and FcεRI expression levels. Our findings support this as multiple cell surface molecules not typically associated with atopy were differentially expressed on the surface of the FIM population. Furthermore, functional assays of sorted monocyte populations also supported differences between populations in antiviral maturation and cytokine release based on surface phenotyping. These data suggested that FIMs are hyper-responsive to both antiviral and IgE-mediated stimulation which may have implications when these cells migrate into tissue during viral infections or allergic airway exacerbations.

In the airway and lung, baseline hyper- or increased inflammatory responses induced by allergic disease in monocytes could result in increased tissue inflammation in the setting of viral infections. Respiratory viruses, particularly RNA viruses that activate pathways like lipo-pIC, are highly associated with allergic asthma exacerbations ^28^. Although none of the lung tissue donors tested positive for respiratory viruses at the time of death (data not shown), within the lung, many of the cell surface markers upregulated on FIMs are associated with pro-inflammatory and antiviral responses. The notable differences between blood and lung monocytes identified in this study begin to characterize the potential changes that occur as these cells move from blood into tissue. For example, CD86, CD40, and ICAM-1 are associated with cellular maturation and ICAM-1 specifically is utilized for vascular and tissue migration. Surface expression of these molecules were lower on blood FIMs compared to the total CD14+ population consistent with these cells potentially in a resting state in the blood. In the lung, these proteins, along with many others, were upregulated on FIMs, consistent with an activated state and altered surface phenotype when transitioning into tissue. This change could imply critical tissue-related functions. For example, MHC II, CD86, and CD80 are all upregulated during viral infections to support antigen presentation and T cell co-stimulation. High expression of these molecules may lead to skewed T cell activation in the setting of either allergen or virus antigen uptake. ICAM-1 was also increased and is the receptor for serogroup A rhinoviruses, thus could modulate how monocytes respond to rhinovirus infections specifically. However, not all the receptors with increased expression are pro-inflammatory. ILT-3 and ILT-4 are immunoglobulin-like receptors and are primarily inhibitory ^29,30^. Similarly, while Axl is an antiviral receptor upregulated by IFN responses, it’s role is to negatively regulate antiviral responses ^31^. We suspect that the combinatorial signals from all of the receptors identified influence the ultimate functional response by FIMs during allergic airway disease and anti-pathogen responses. The increased proportion of FIMs, in the blood and subsequently the lung, combined with altered functions may ultimately lead to dysregulation of cellular responses – both in the FIM population and via paracrine responses by other monocytes or immune cells, and other target cells in the lung milieu.

This study provides new insights into how atopic disease influences development and differentiation of monocyte subsets. To our knowledge this is the first study to identify unique cell surface phenotypes and functional responses in monocytes with high levels of both the FcεRI receptor and surface-bound IgE. Our use of high dimensional flow cytometry determined how allergic disease and IgE levels are related to expression of many monocyte surface proteins not typically associated with allergic disease. Our results are consistent with other studies including multiple recent studies using RNA sequencing approaches which have also identified multiple transcriptionally distinct monocyte subsets ^8,32^. Some of these studies also included airway samples from subjects with allergic asthma ^8^. However, these studies did not specifically identify IgE-bound populations, which are not identified transcriptionally as IgE is bound from the serum. Our approach specifically detected differences related to surface-bound IgE and it is clear from these results that IgE receptor expression and IgE binding – whether via the high (FcεRI) or low (CD23) affinity receptors – are not equivocal. We were able to induce IgE-driven effects in the low IgE-bound population, suggesting that, despite low detection, some receptors are occupied and functional. Future studies will build on these findings to specifically evaluate how IgE-bound monocytes differ in broader functionality and transcriptionally from unbound (or low-bound) monocytes in the context of allergic inflammation. We also identified several subjects with discordant allergic history and serum IgE levels, either high IgE and no allergic history or low IgE with allergic history. Although our allergic history was not confirmed with allergen skin testing, subjects who reported no allergic history but had high serum IgE had cellular phenotypes similar to those with both allergy history and high serum IgE levels. Further comparison of these discordant individuals could shed critical information on how allergen tolerance is established in the setting of high IgE levels.

Finally, using a unique lung tissue biorepository, we were demonstrated that the monocyte phenotypes associated with atopy we identified in the blood are also present in lung tissue from individuals who died of lethal asthma exacerbations. This provides a foundation for future work to further evaluate both similarities and differences between blood and lung monocytes to identify which how monocytes transition from blood into tissue during allergic airway inflammation and respiratory virus infections. This work has expanded our knowledge of how allergic disease regulates monocyte phenotypes and functions in both blood and lung monocytes with FIMs potentially serving as an additional biomarker of atopy.

## Materials and methods

### Human Subjects

Primary human immune cells were isolated from 2 sources: leukocyte-enriched blood samples from unknown subjects acquired from blood banks and peripheral blood samples from healthy and allergic subjects (Table 1). Unknown subjects were deemed healthy enough for routine blood donation by the blood bank, with limited demographic information (age and sex) available. In a select subset of these unknown subjects, we were able to obtain serum to evaluate IgE levels, otherwise no clinical information (*i.e.* atopic status or allergic history) was obtained. For all known subjects (Table 1), written informed consent and assent were obtained. Known subjects were recruited from the local Rochester community and completed a medical and allergy history questionnaire. Peripheral blood was obtained to isolate PBMCs and serum was collected to measure IgE levels. Studies were approved by the University of Rochester Research Studies Review Board (RSRB, Study #00006081).

### Serum IgE levels

Serum was isolated from 2-5mL of peripheral blood by serum separator tubes by spinning at 1200 x g for 10 minutes. Serum was harvested and frozen in 0.5mL aliquots at −80°C until analysis. To measure serum IgE levels, frozen serum was sent to the University of Rochester Medical Center Clinical Microbiology Laboratory and IgE levels were measured via a clinically validated test. Levels were reported in international units (IU) per mL. Subjects were divided into high IgE (>100IU/mL) or low IgE (<100IU/mL) groups, and by allergic history as shown in Table 1 for data analysis purposes.

### Reagents and Media

Phosphate Buffered Saline (PBS) with fetal bovine serum (FBS) and EDTA (complete PBS, cPBS) and complete Roswell Park Memorial Institute Medium 1640 (cRPMI) were prepared as previously described ^3,4,27^. All medias were purchased from Gibco unless otherwise stated and FBS was purchased from Biowest (Riverside, MO), and Ficoll-Paque™ PLUS (Cytiva, Uppsala, Sweden).

### Preservation of PBMCs from Blood

PBMCs were isolated via Ficoll gradient centrifugation. PBMCs were preserved frozen at 1×10^7^ cells per mL in freezing medium (10% DMSO, 50% FBS, 40% cRPMI). Cells were slow cooled at −80°C overnight in a Mr. Frosty^TM^ container (Thermo Scientific), then transferred to liquid nitrogen for long term storage. For analysis, PBMCs were quick thawed in a 37°C water bath for 2-3 minutes then washed once in cRPMI at 4 times the volume, pelleted and washed once more in cPBS. PBMCs were counted for viability with trypan blue (typically >90%) and processed for various downstream analyses.

### Purification and Culture of Monocytes from PBMCs

To isolate monocytes, thawed PBMCs were resuspended at up to 5×10^7^ cells/mL in cPBS. Monocytes were purified by negative selection as previously described ^2–4,27^ using the EasySep^TM^ Human Monocyte Enrichment kit (catalog #19059), per manufacturer’s instructions (STEMCELL Technologies, Vancouver). Isolated monocytes were then either cultured directly or stained for flow cytometry sorting. Monocytes were then cultured in complete RPMI medium (cRPMI) at a concentration of 1 × 10^6^ cells/ml in 96-well flat bottom plates. Cells were stimulated immediately after purification or sorting to ensure adequate crosslinking of native surface bound IgE and minimize downregulation of surface receptors. IgE-mediated allergic stimulation was performed by crosslinking cell surface bound IgE using a goat anti-human IgE polyclonal antibody (αIgE 10 μg/ml), with goat IgG isotype control (10 μg/ml) or media alone used as controls ^2^. To activate antiviral cellular responses, cells were treated with liposomal complexed low molecular weight poly(I:C). Liposomal poly(I:C) complexes were made by mixing poly(I:C) with Lipofectamine 2000 at a ratio of 1µg of poly(I:C) to 1µl of Lipofectamine 2000 reagent in RPMI media. The mixture was incubated at room temperature for 30 minutes and added to monocyte cultures at a concentration of 1µg/mL (based on poly(I:C) concentration). Cells were cultured at 37°C for 24 hours post stimulation and supernatants harvested for ELISA analysis and cells isolated for flow cytometry.

### Flow Cytometry Analysis of Surface Antigens

PBMCs were analyzed with a multiplex antibody panel using the fluorochrome-conjugated anti-human antibodies listed in Supplemental Table 1. Optimization of antibody concentrations and the use of FMO controls were performed during the panel design process. Clones were chosen based on unique specificities including: 1) AER-37 (CRA-1) which does not compete with IgE binding and thus detects total FcεRIα receptor levels ^33^, 2) the anti-human IgE clone chosen detects receptor bound IgE, and 3) CD14 and CD16 were non-competing with the clones in the lineage cocktail. Staining and flow cytometry were performed as previously described ^2^. Samples were acquired on a Cytek Aurora spectral cytometer (Cytek Biosciences, Fremont, CA). Live spectral unmixing was performed during acquisition by the Cytek SpectroFlo software using single fluorochrome stained beads for each antibody (UltraComp Compensation Beads, ThermoFisher Scientific), a live/dead single stain cell control, and an autofluorescence extraction using unstained cells (PBMCs or monocytes). Unmixed data was analyzed using FCS express version 8 or the FlowJo version 10 software (FLOWJO, LLC, Ashland, OR).

For cell sorting, purified monocytes were similarly stained for viability (live dead blue) followed by surface staining with a focused panel (CD14, FcεRIα, IgE, CD16) and then sorted on the Cytek AS spectral sorting cytometer. Live CD14+ cells were sorted into two populations: FcεRIα+IgE+ (FIMs) and the remaining CD14+ monocytes (Not Gate). FMO controls for FcεRIα and IgE antibodies were used to identify the FcεRIα+IgE+ positive versus “not gates”.

High dimensional analysis of flow cytometry data was performed as follows using FlowJo version 10 software and R studio: Individual .fcs data files were cleaned (FSC, SSC, and time gates) and gated on live single cells. Categorical metadata was added to each file including subject number, batch, serum IgE category, allergic disease, and treatment if indicated. Batch correction of live single gates was performed using FlowJo10 and R using the CytoNorm ^34^ plugin in FlowJo. Batch correction was performed using internal identical “control” PBMC samples, from discarded leukocyte fractions which represented all cell populations found in both low and high IgE subjects. Normalized batch corrected files were then used for downstream high dimensional analysis. In FlowJo, batch corrected files were gated to the target populations of interest: monocytes (CD14+) and DCs and basophils (Lin-), or only CD14+ monocytes. This population was concatenated and dimensionality reduction performed using UMAP or PHATE plugins (Supp. Fig. 1G). Traditional gating strategies were used to also identify monocyte, DC, and basophil populations (Supp. Fig. 1), which were overlaid on UMAP or PHATE maps. For experiments using stimulated purified monocytes, batch correction was performed using a representative population (10,000 cells) from each sample and concatenating as the control sample representative of all samples, subjects, and cell populations across the batch. This combined control sample was then used for batch correction using CytoNorm.

### Flow cytometry of monocytes in lung tissue

Cryopreserved dissociated lung tissue was obtained from the BioRepository for Investigation of Diseases of the Lung (BRINDL). The BRINDL repository contains dissociated lung tissue samples from >500 lungs obtained post-mortem, and includes basic demographic, clinical, and histopathology information^35,36^. We selected samples from adult donors either with a known history of asthma (n=7) or no known history of asthma or lung disease (n=7), matched for age and sex as able. Six of the seven individuals with asthma died of lethal asthma exacerbation (Supp. Table 2) and histopathology information for the asthma tissue all had findings consistent with allergic inflammation. Dissociated lung tissue was thawed similar to the PBMC thawing protocol, counted for viability with trypan blue, and processed for flow cytometry. Anti-human CD45 antibody was added to the flow cytometry panel (Supp. Table 1) to identify immune cells in the heterogeneous lung samples.

### Cytokine Secretion Analysis

Supernatants were stored at −80°C until use. The following ELISA kits were used according to manufacturer recommendations: ELISA Max human ELISA for IL-6, IL-10, or CXCL10 (IP-10) (Biolegend, San Diego, CA), Absorbance was measured on SpectraMax iD3 microplate reader (Molecular Devices, San Jose, CA) per manufacturer’s instructions.

### Data Analysis and Statistics

The number of biological replicates for a given experiment are noted in the figure legends. Data in bar graphs are presented as the mean with standard error, with individual biological replicates overlaid. Other data are presented as means ± standard error of the mean (SEM) or standard deviation (SD) as noted. In experiments containing 3 or more conditions and comparisons, ANOVA with mixed effects analysis followed by post hoc comparisons were performed using Sidak correction for multiple comparisons performed when appropriate. For experiments comparing 2 conditions, paired two-tailed t tests were performed with Welch’s correction. Mann-Whitney U test was used for comparison of subjects and variables for categorical variables, high vs. low IgE or allergic vs. no allergic history. For continuous variable analysis, linear regression analysis with two-tailed t-test was performed and Pearson’s coefficients (R^2^) and p values determined. p<0.05 was considered significant, pertinent p values are denoted as follows: *p<0.05, **p<0.01, and ***p<0.001, or ns (non-significant) for p>0.05. All statistical analyses were performed using GraphPad Prism version 10.

## Acknowledgements

We thank the participants, without whom this study would not be possible. We want to thank the URMC Vaccine Treatment and Evaluation Unit (VTEU), Dr. Jennifer Nayak, Dominique Williams, and Michelle Heiman (Division of Pediatric Infectious Disease, URMC) for assistance with study subject recruitment and study visit completion, and Nelson Huertes for technical assistance. We acknowledge the LungMAP Human Tissue Core (funded through NIH NHLBI U01-HL122700 and U01-HL144861), specifically Heidie Huyuck and Dr. Gloria Pryhuber, for providing the lung tissue samples. We acknowledge the staff of the University of Rochester Medical Center Flow Cytometry Core for assistance with flow cytometry analysis. We thank the following funding sources: NIH NIAID, 5K08AI163380 (RKR), URMC VTEU Pilot Grant Program, and the Department of Pediatrics, University of Rochester Medical Center.

## Abbreviations

FIM: FcεRIα+IgE+ monocyte
αIgE: Goat anti-human IgE antibody, IgE crosslinking condition
CD: cluster differentiation
DC: dendritic cell
FcεRI: high-affinity IgE receptor
FcεRIα: alpha subunit of the high-affinity IgE receptor
IgE: immunoglobulin E
IgG: Goat whole IgG isotype control antibody
IL: interleukin
IFN: interferon
PBMCs: peripheral blood mononuclear cells

