## Supplemental Materials for "FcεRI+IgE+ monocytes are linked to atopy and allergic inflammation with distinct phenotypes and enhanced antiviral responses"

| Supplemental Table 1: Spectral Flow Cytometry Panel | | | | | |
|---|---|---|---|---|---|
| **Antigen** | **Fluor** | **Dilution** | **Clone** | **Company** | **Catalog #** |
| Viability Stain | Live Dead Blue | 1/1000 | n/a | ThermoFisher | L23105 |
| IgE | BUV395 | 1/50 | G7-26 | BD Biosciences | 744321 |
| CD40 | BUV496 | 1/50 | 5C3 | BD Biosciences | 741159 |
| CD23 | BUV563 | 1/50 | EBVCS-5 | BD Biosciences | 749450 |
| CD14* | BUV737 | 1/50 | MφP9 | BD Biosciences | 741897 |
| CD86 | BUV805 | 1/50 | 2331 (FUN-1) | BD Biosciences | 742032 |
| CD80 | BV421 | 1/50 | L307.4 | BD Biosciences | 564160 |
| (ICAM1) = CD54 | Pacific Blue | 1/100 | HA58 | Biolegend | 353110 |
| Siglec-6 | BV480 | 1/50 | 767329 | BD Biosciences | 747914 |
| CD123 | BV605 | 1/100 | 6H6 | Biolegend | 306026 |
| HLA-DR (MHC II) | BV615 | 1/400 | L243 | BD Biosciences | 753690 |
| CD1c | BV650 | 1/100 | F10/21A3 | BD Biosciences | 742749 |
| CD11c | BV711 | 1/100 | B-ly6 | BD Biosciences | 563130 |
| ILT-3 (CD85k) | BV786 | 1/50 | ZM3.8 | BD Biosciences | 742810 |
| ILT-4 (CD85d) | PerCP-eFluor 710 | 1/50 | 42D1 | Fisher/Invitrogen | 50-245-903/ 46514941 |
| Lineage (CD3, CD19*, CD20*, CD16*, CD14*, CD56) | FITC | 1/40 | UCHT1; HCD14; 3G8; HIB19; 2H7; HCD56 | Biolegend | 348801 |
| OX40L (CD252) | PE | 1/50 | 11C3.1 | Biolegend | 326308 |
| FcεRIα | PE-Dazzle594 | 1/50 | AER-37 (CRA-1) | Biolegend | 334634 |
| CD141 | PE-Cy7 | 1/800 | M80 | Biolegend | 344110 |
| Clec9A (CD370) | AF647 | 1/100 | 3A4/Clec9A | BD Biosciences | 564267 |
| CD16* | AF700 | 1/100 | B73.1 | Biolegend | 360718 |
| CD45[#] | APC/Fire810 | 1/100 | HI30 | Biolegend | 304076 |

*Non-competing clones used
[#]Used only for lung tissue samples

| Supplemental Table 2: Lung Tissue Description | | | | |
|---|---|---|---|---|
| | **Asthma History** | **Age** | **Sex** | **Cause of Death** |
| 1 | No | 40y | F | CVA/Intracranial Hemorrhage |
| 2 | No | 29y | F | Head Trauma/GSW |
| 3 | No | 19y | M | Anoxia/Drug Intoxication |
| 4 | No | 24y | M | Accidental Head Injury/Motor Vehicle Accident |
| 5 | No | 25y | F | Drug Intoxication |
| 6 | No | 59y | M | CVA/Intracranial Hemorrhage |
| 7 | No | 20y | F | Head Trauma/Motor Vehicle Accident |
| 8 | Yes | 24y | M | Anoxia/Cardiopulmonary Event |
| 9 | Yes | 23y | F | Acute Fatal Asthma |
| 10 | Yes | 21y | M | Head trauma/GSW |
| 11 | Yes | 24y | F | Asthma/Status Asthmaticus |
| 12 | Yes | 39y | F | Acute Asthma/Drug intoxication |
| 13 | Yes | 25y | F | Asthma/Acute Asthma Exacerbation |
| 14 | Yes | 22y | F | Asthma Exacerbation |

### Supplemental Figure 1

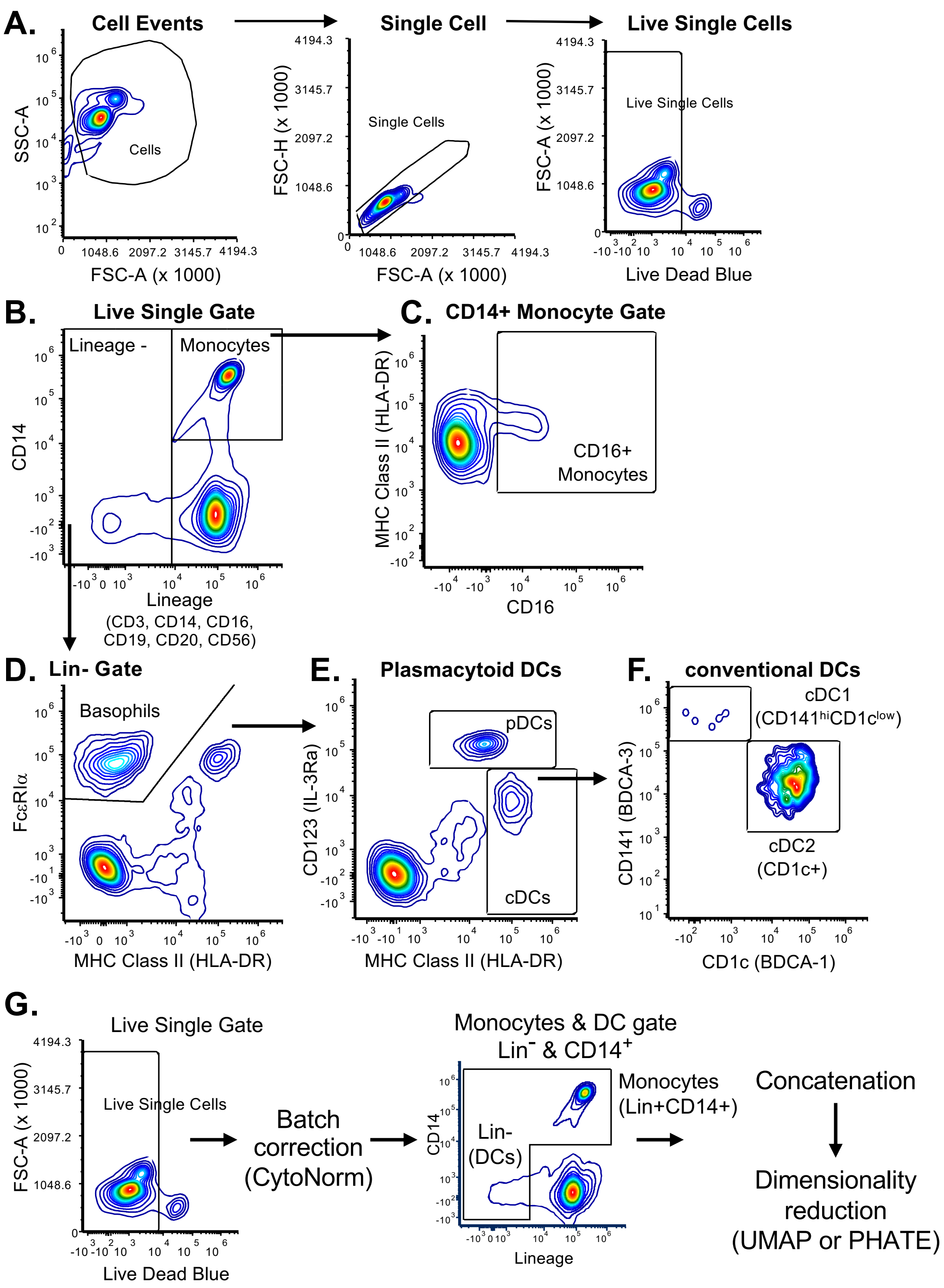


**Supplemental Figure 1:** Flow cytometry gating and analysis strategy. Cells were first gated to live single cell events (A). (B) Total monocytes were identified as Lineage+CD14+ and (C) then CD16+ subset downstream. (B) Lineage negative population was used to identify (D) basophils (Lin-FcεRIα+, MHC II neg, also confirmed as CD123 high in data not shown), (E) plasmacytoid dendritic cells (Lin-, CD123hi, MHC II hi) and conventional (myeloid) dendritic cells (Lin-CD123lo, MHC II hi), then into (F) cDC1 (CD141hi) and cDC2 (CD1c+) subsets. (G) Gating and analysis strategy shown for dimensionality reductions UMAP and PHATE.

### Supplemental Figure 2

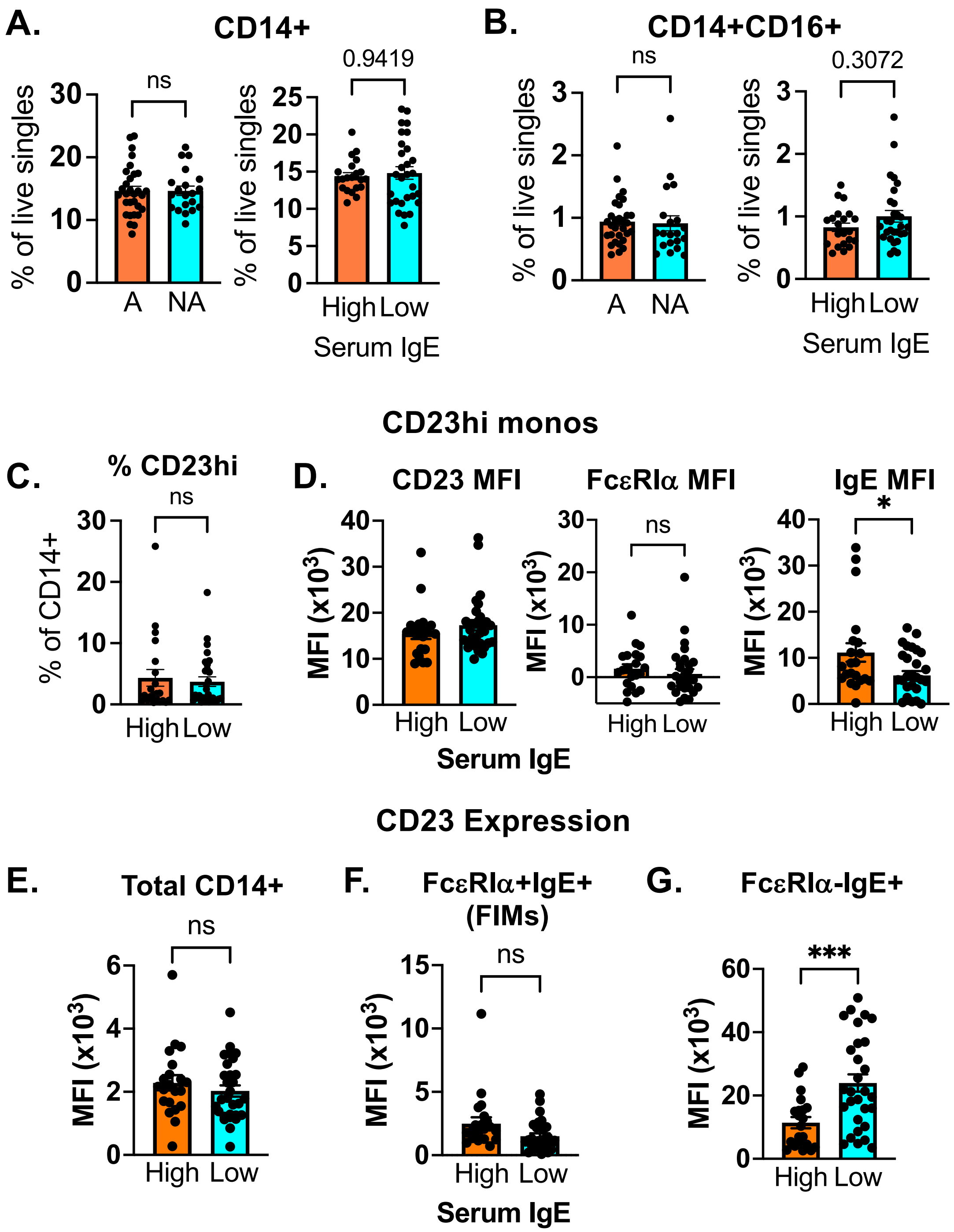


**Supplemental Figure 2:** Comparison of proportions and surface molecule expression on other monocytes populations (CD14+, CD14+CD16+, and CD23hi) based on allergic history and serum IgE.

Supplemental Figure 3

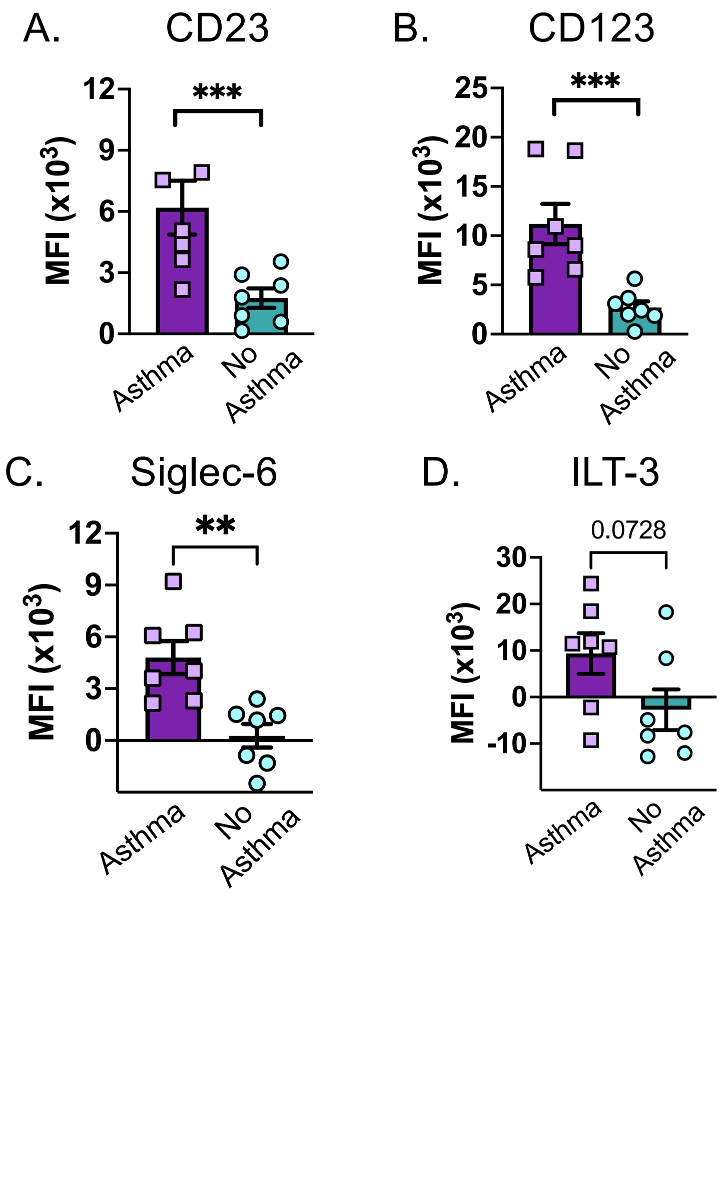


**Supplemental Figure 3:** Comparison of select markers on CD14+ monocytes in lung tissue from subjects with asthma or no asthma history. Unpaired t-test with Welch's correction; *$p<0.05$, **$p<0.01$, ***$p<0.001$, and not significant (ns)
